# Structural heterogeneity of amyloid aggregates identified by spatially resolved nanoscale infrared spectroscopy

**DOI:** 10.1101/2022.05.07.491036

**Authors:** Siddhartha Banerjee, Brooke Holcombe, Sydney Ringold, Abigail Foes, Ayanjeet Ghosh

## Abstract

Amyloid plaques, composed of aggregates of the amyloid beta (Aβ) protein, are one of the central manifestations of Alzheimer’s disease pathology. Aggregation of Aβ from amorphous oligomeric species to mature fibrils has been extensively studied. However, significantly less in known about early-stage aggregates compared to fibrils. In particular, structural heterogeneities in prefibrillar species, and how that affects the structure of later stage aggregates are not yet well understood. Conventional spectroscopies cannot attribute structural facets to specific aggregates due to lack of spatial resolution, and hence aggregates at any stage of aggregation must be viewed as having the same average structure. The integration of infrared spectroscopy with Atomic Force Microscopy (AFM-IR) allows for identifying the signatures of individual nanoscale aggregates by spatially resolving spectra. In this report, we use AFM-IR to demonstrate that amyloid oligomers exhibit significant structural variations as evidenced in their infrared spectra, ranging from ordered beta structure to disordered conformations with predominant random coil and beta turns. This heterogeneity is transmitted to and retained in protofibrils and fibrils. We show for the first time that amyloid fibrils do not always conform to their putative ordered structure and structurally different domains can exist in the same fibril. We further show the implications of these results in amyloid plaques in Alzheimer’s tissue using infrared imaging, where these structural heterogeneities manifest themselves as lack of expected beta sheet structure.

## Introduction

Aggregation of the amyloid beta (Aβ) protein into insoluble fibrillar deposits or plaques is a pathological hallmark of Alzheimer’s disease (AD)^1-2^. The mere presence of amyloid plaques, however, does not show correlation with cognitive decline^1, 3-4^, and recent years have seen a paradigm shift in the pathological implications of Aβ plaques. It is now believed that soluble Aβ oligomers and prefibrillar aggregates and not mature fibrils are the primary neurotoxic species in AD^5-8^. The secondary structure of Aβ in fibrils and prefibrillar aggerates has also been studied extensively, and it is known that mature Aβ fibrils have a parallel beta sheet secondary structure^9-13^, but Aβ oligomers may have antiparallel beta sheet structure^14-15^. Although these studies provide valuable information, isolation and structural analysis of the amyloid aggregates formed at different stages of the aggregation still remains a challenge. The variations in protein structure within one subgroup of aggregates corresponding to a specific time point of aggregation are yet to be well understood. This is particularly relevant for early-stage aggregates such as oligomers and protofibrils. It is well-known that mature amyloid fibrils exhibit polymorphism: fibrils can have different morphologies which not only correspond to differences in molecular structure^12, 16-17^, but are also associated with different stages of AD progression^11, 18^. However, little is known about structural polymorphism of early-stage aggregates. Essentially, it is not known if a particular type of aggregate can exhibit different secondary structures and if/how such structural heterogeneity can influence later stage aggregates. This gap in knowledge stems in part from the lack of suitable techniques that offer structural sensitivity and simultaneously are capable of spatially resolving specific aggregate species. Conventional approaches for protein structure determination, such as Nuclear Magnetic Resonance (NMR) and infrared spectroscopies do not offer spatial resolution and provide average spectra. As a result, it is not possible to assign spectral and hence structural attributes to specific aggregates, and it has to be assumed that all aggregates at a given stage of aggregation have the same average structure. The development of cryo-Electron Microscopy (cryo EM) has been able to address this limitation by spatially resolving structural variations in amyloid fibrils and promises to be an invaluable tool towards understanding amyloid structure^19-22^. However, the applications of cryo-EM have been limited largely to mature fibrils, and hence there is need for experimental approaches that can isolate the structural facets of prefibrillar and early-stage aggregates. Integration of infrared spectroscopy with Atomic Force Spectroscopy (AFM) presents a perfect solution to this challenge. AFM is a technique routinely used to study amyloid aggregates, and AFM-IR expands its capabilities by allowing for measurement of nanoscale infrared spectra of individual aggregates by utilizing the photothermal effect, where the local thermal expansion resulting from infrared absorption by a sample is sensed by an AFM tip^23-28^ (Fig. 1). AFM-IR thus combines the structural sensitivity of infrared spectroscopy with the nanoscale spatial resolution of AFM and allows for the identification of structural heterogeneity of aggregates by acquiring spectra of each member of the ensemble. In this study, we use AFM-IR to probe the structural distribution of Aβ42 oligomers and early-stage fibrils. Our findings show for the very first time that Aβ oligomers can exhibit a range of secondary structures and the integration of these heterogeneous oligomers into fibrils leads to fibrillar structural disorder, wherein different regions of the same fibril exhibit different structure, demonstrating that amyloid fibrils can deviate significantly from their putative structural uniformity.

**Figure 1.**
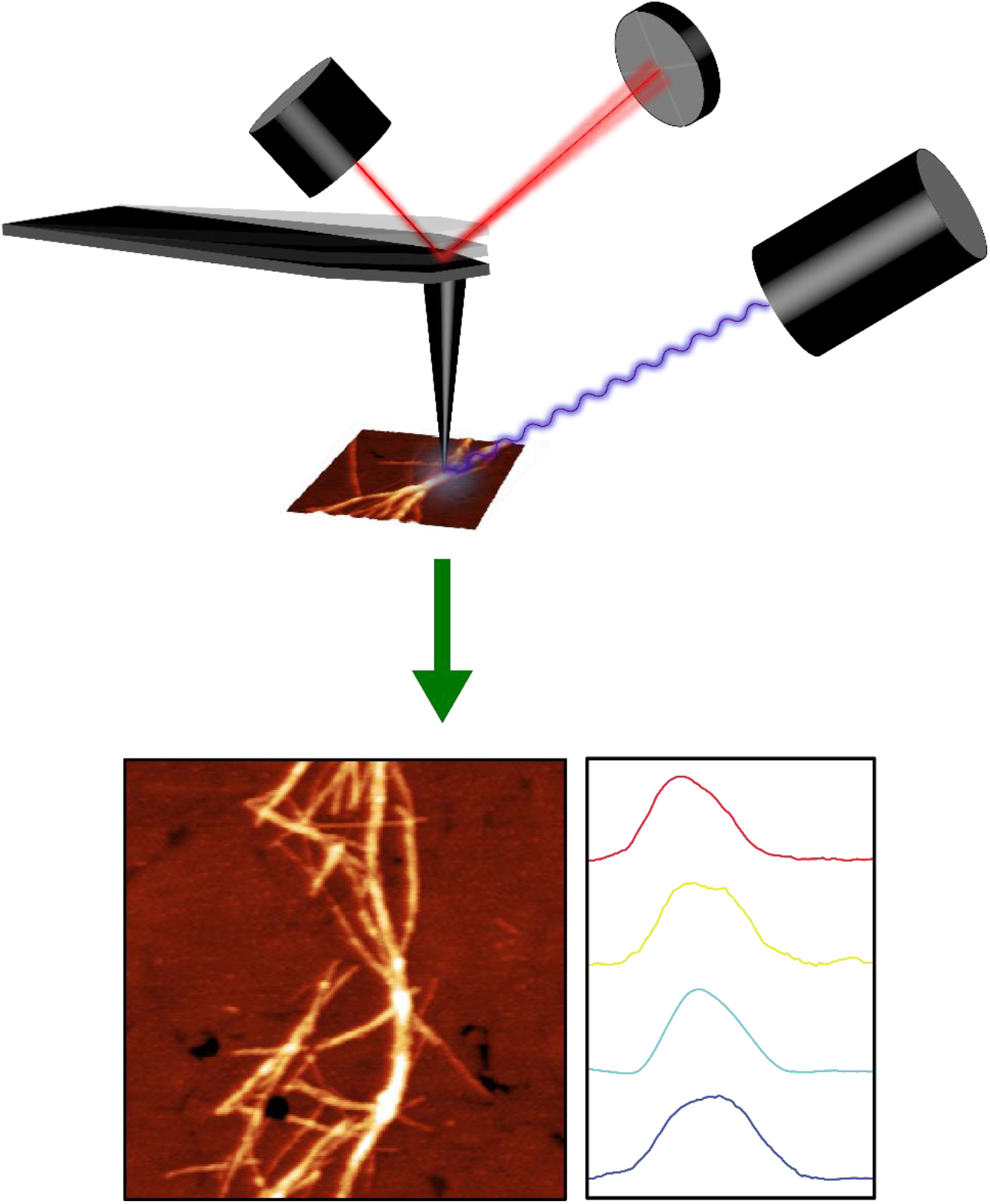
Schematic description of AFM-IR. The AFM-IR measurement involves resonant excitation of the sample with infrared radiation, which leads to thermal expansion. The amplitude AFM cantilever oscillations resulting from this expansion are proportional to the infrared absorbance and lead to spatially localized nanoscale infrared spectra.

## Results

### AFM-IR reveals heterogeneity in secondary structure of prefibrillar aggregates

The aggregation of Aβ42 was monitored by depositing the aggregation mixture at different time intervals and acquiring AFM images. The experimental methods are detailed in the Methods section. Oligomers, protofibrils and fibrils were observed at different stages of aggregation. Fig. 2A shows the AFM topographic image of Aβ42 oligomers observed after 3 days. Single isolated individual oligomers can be visualized. In addition to oligomers, elongated protofibrillar aggregates are also observed alongside oligomers, which are discussed later. The observed oligomeric species are globular or spherical in morphology; the height of these globular aggregates was measured to be 2.6±0.6 nm (Suppl Fig. 1). Representative IR spectra recorded from these oligomeric species are shown in Fig 2B; additional spectra are shown in the Supplementary Information (Suppl Fig. 2A). The amide I absorption band exhibits significant heterogeneities between different oligomers and a single spectral type cannot be attributed to the oligomeric aggregates. The amide I band shows distinct peaks at 1634 cm^-1^ and 1666 cm^-1^. It is well known that amide-I infrared spectra are reflective of protein secondary structure, and thus the observation of different spectral types for oligomers clearly demonstrates that despite no significant morphological differences, early-stage oligomeric aggregates have significant variations in secondary structure. Interpreting the structural implications of the peaks observed and understanding the relative distributions of secondary structures contributing to the spectra requires spectral deconvolution, which we discuss later.

**Figure 2.**
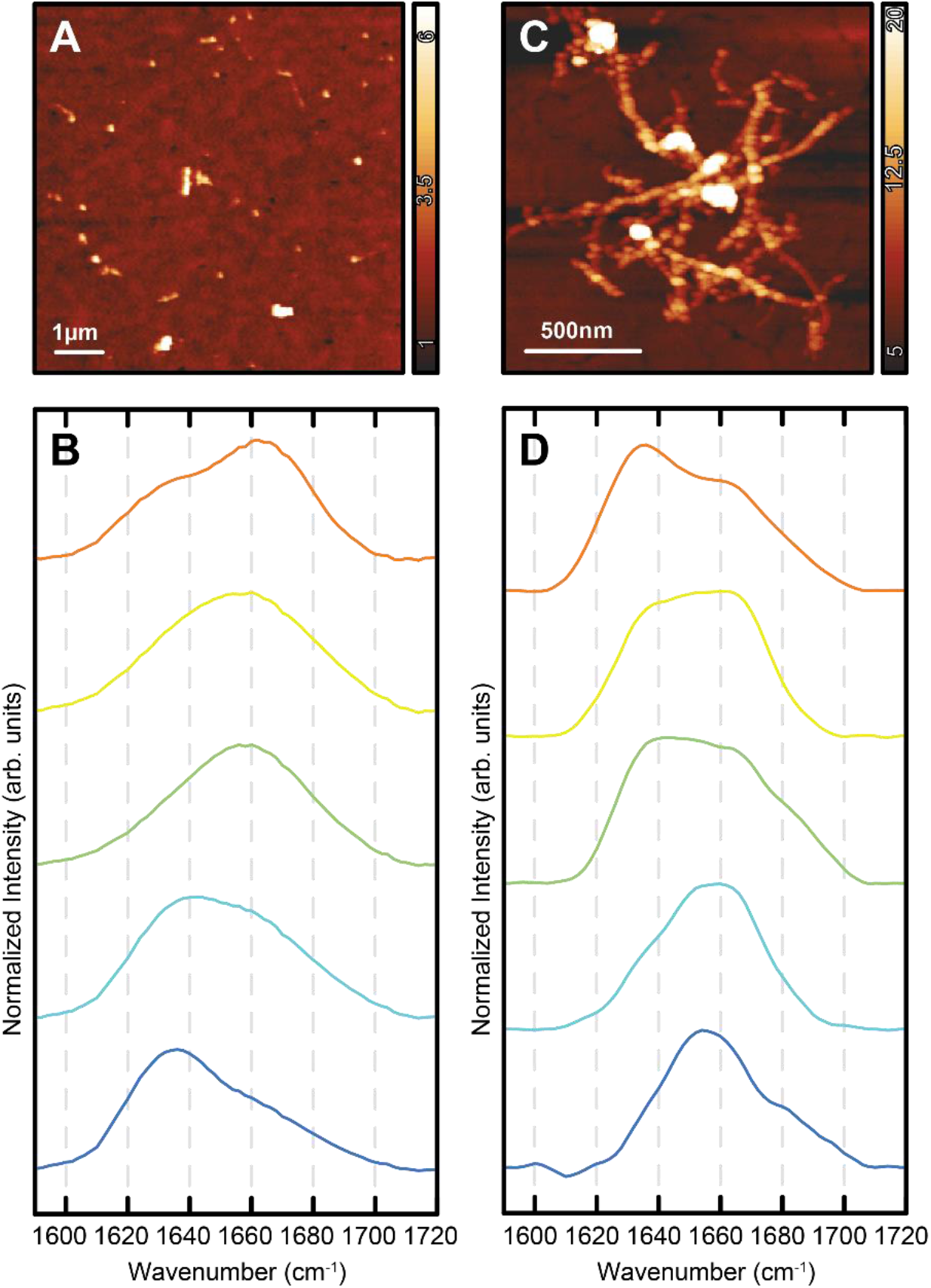
Structural characterization of individual early-stage Aβ42 aggregates. (A) AFM topographic image of Aβ42 oligomers generated after 3 days of incubation in PBS buffer. (B) Representative IR spectra of the amide I region obtained from different individual oligomers. (C) AFM image of Aβ42 protofibril and (D) corresponding IR spectra recorded from different regions of the protofibril cluster.

Protofibrillar aggregates were observed alongside oligomers. The morphology of a single protofibrillar aggregate is shown in Fig. 2C. The protofibril appears significantly different from the usual amyloid fibril morphology. The protofibrils observed do not appear to be smooth, rod-like amyloid fibrils, rather they are composed of individual globular aggregates attached with each other along the long axis of the protofibrils (Fig. 2C). This feature is more evident when the topographic image is shown in a rainbow color scheme (Suppl Fig. 3A), and also in the cross-sectional profile along the long axis (Suppl Fig. 3B), which exhibits multiple peaks, and strongly indicates the presence of individual oligomeric features in the composition of protofibrils. IR spectra were recorded at arbitrarily chosen points on different protofibrils, and also along the length of the protofibrillar aggregate shown in Fig. 2C. Representative spectra are shown in Fig. 2D. It is evident that the amide I absorption bands shows distinct differences between different protofibrils (Suppl Fig. 2B) and also within a single protofibril. The spectral variations are consistent with those observed for oligomers: the spectra exhibit a peak around ∼1632 cm^-1^ with a second shoulder peak at ∼ 1666 cm^-1^. In some spectra, the intensities of these bands are nearly equal, leading to a plateau like appearance for the spectra from 1630-1670 cm^-1^. These spectral heterogeneities indicate the presence of varied secondary structures of Aβ42 in protofibrils. The spectral similarities between oligomeric species and protofibrils strongly suggest that the protofibril formation involves integration of individual oligomers which retain their structural facets even after incorporation into the protofibrillar structure.

### Amyloid fibrils can exhibit heterogeneous structure3

To understand the effect of this heterogeneity on later stage aggregates, we investigated the structural distribution of fibrils. Amyloid fibrils were mostly observed after 7 days of incubation. Fig. 3A shows the AFM topographic image of a representative fibril. The fibril morphology is distinctly different compared to the protofibrils and no distinct globular bead like features along the fibril axis are detected. The height of the fibril, as measured at different regions of the fibril, is plotted as a histogram in Suppl Fig. 4. The mean height is 11.5±5.2 nm which is significantly higher compared to the protofibrils. Additional fibril images are also shown in Suppl Fig. 4. Similar to the protofibrils, IR spectra were acquired at multiple points along the fibril axis and representative spectra are shown in Fig. 3B. Additional spectra are shown in Suppl Fig. 2. The spectra exhibit similar heterogeneity observed in earlier aggregates with peaks at ∼ 1660 cm^-1^, ∼ 1630 cm^-1^ or both. We note that some fibrillar spectra exhibit a distinct difference from those described above and show a broad amide I absorption centered at ∼1668cm^-1^ or even a distinct peak at ∼1680cm^-1^, which suggests the possibility of a different secondary structure in fibrils absent in earlier oligomers.

**Figure 3.**
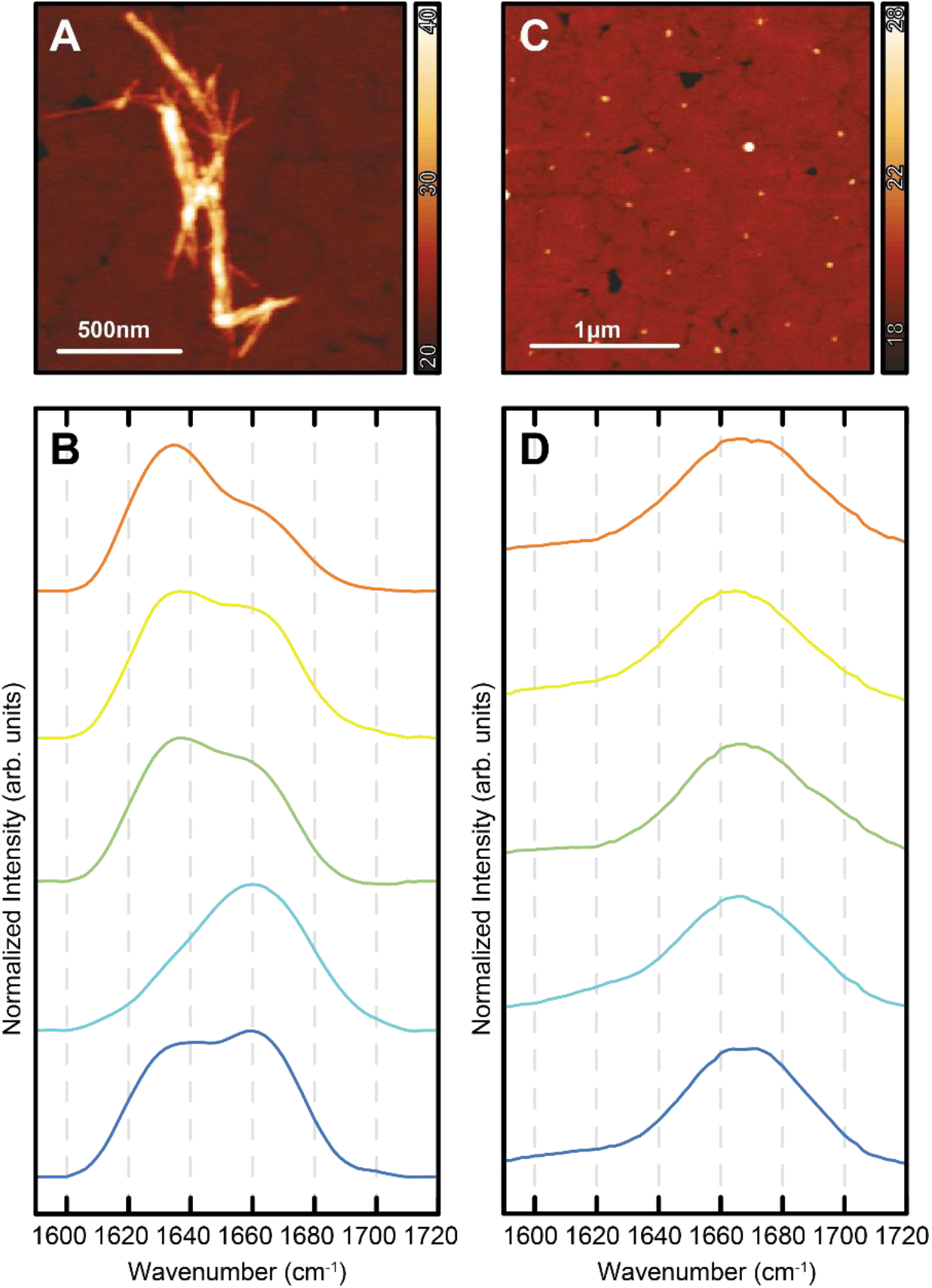
Structural characterization of Aβ42 fibrils and later-stage oligomers. (A) AFM image of the Aβ42 fibril on gold substrate. (B) IR spectra obtained from different points along the fibril long axis. (C) AFM image of the Aβ42 oligomers observed alongside fibrils. (B) Representative IR spectra recorded from different oligomers.

### Later stage oligomers are structurally homogeneous

In addition to fibrils, we have also observed a significant number of oligomers after 7 days of aggregation, as shown in the AFM topographic image (Fig. 3C). The height value of these oligomers has been measured to be 3.1 ±0.7 nm (Suppl Fig. 5). IR spectra have been recorded on these individual oligomers and shown in Fig. 3D and Suppl Fig. 2. Interestingly, the amide I absorption band does not show heterogeneity, as opposed to the spectra observed for the oligomers generated before the fibril formation. The peak of the absorption band is centered around ∼ 1668 cm^-1^, similar to some of the spectra observed in fibrils, indicating that these oligomers can be participating in fibril formation. Interestingly, these oligomers do not exhibit the peak around ∼1630 cm^-1^, indicating a distinct deviation from 3-day oligomeric structure, which could potentially hinder their participation in further fibril formation, consistent with their abundance at later stages of aggregation.

### Structural implications of spectral heterogeneity

These results unequivocally demonstrate that early-stage Aβ42 aggregates are polymorphic in their structure, and the structural diversity in oligomers is retained and propagated to protofibrils and fibrils. This is a particularly surprising observation, since it is expected that the aggregation mechanism involves a transition from more disordered oligomeric species to ordered fibrils^9, 29-32^. Our observations indicate that fibrils may not be necessarily having the ordered structures they are envisioned to be, and the structural variations and/or disorder of early-stage aggregates can still persist in fibrils. It is known from solid state NMR and more recent cryo-EM studies that a part of the polypeptide sequence can exist in a disordered state even in highly ordered amyloid fibrils^33-35^. In fact, spectral line broadening in infrared measurements of amyloid species have been observed and attributed to structural disorder^30, 32, 35^. However, in all cases, the structural disorder has been assumed to arise only from different degrees of ordering within the amyloid peptide, and not from coexisting physical domains exhibiting variations in secondary structure. To the best of our knowledge, heterogeneities in secondary structure of beta amyloid aggregates that are not reflected in morphology have not been identified before. Previous AFM-IR studies on α-synuclein aggregates have revealed variations in secondary structure between spherical and elongated, ‘fibril-like’ oligomers^36^. Our results demonstrate for the first time that structural variations can arise from different oligomeric aggregates which in turn lead to spatial variations of secondary structure along a single amyloid fibril. Interestingly, we have previously identified structural heterogeneities in tau fibrils, wherein fibrils exhibiting different spectra were identified, but no significant spectral variations were observed within the same fibril^37^. The results reported here point towards a fundamental difference in the structure of early-stage aggregates of beta amyloid and tau. AFM-IR has been used to investigate amyloid aggregates before^38-39^, but spectral variations along fibrils or heterogeneities observed herein have not been reported. It must be noted that the fibrillar structures investigated in this report are not mature fibrils, and the observations reported herein pertain to early stage fibrillar aggregates. It is possible that mature fibrils may exhibit less pronounced structural variations. Hence potential effect of aggregation conditions and/or time on the structural variabilities cannot be ruled out.

The spectra, as described above, clearly indicate the presence of structural variations in early-stage amyloid aggregates. However, amide I spectra are usually convolutions of overlapping bands from different conformations, and hence the structural underpinnings of the AFM-IR spectra cannot be readily determined without spectral deconvolution. To get insights into the relative populations of secondary structures contributing to the spectra, we adopt a spectral deconvolution approach that utilizes non-negative matrix factorization (NMF)^40-42^. The spectra of all the aggregates were deconvoluted together with NMF, which essentially makes this approach similar to global spectral fitting and allows for interpretation of spectral differences in terms of relative population of the same underlying sub-states. The details of the NMF analysis are provided in the Methods section. We find that 3 components/sub-bands adequately describe the spectral variations which are shown in Fig. 4A. Component 1 has a peak centered at 1632cm^-1^ and a weaker shoulder at 1660cm^-1^. These features are representative of typical ordered parallel beta sheet spectra, as recorded for amyloid aggregates with FTIR^30-32^. The second component is primarily comprised of a broad peak at 1658cm^-1^. It is known that disordered random coil structures absorb at ∼1654cm^- 1^, whereas beta turns can exhibit bands typically at 1660cm^-1^ and higher^43^. Component 2 very likely represents contributions from both random coils and turns; however further deconvolution of this band can be challenging without a priori structural information. However, it should also be noted in this context that presence of significant populations of either of these structures represents a deviation from the ordered fibril structure; hence component 2 can be thought of representative of fibrillar structural disorder. The 3rd component exhibits a prominent peak at 1682cm^-1^, which is typically attributed to antiparallel beta sheets^14, 31, 36^. A weaker peak is observed at 1610cm^-1^, which is too low to be attributable to typical beta sheet structure. However, we note that Lomont and coworkers^44^ have reported the presence highly ordered beta sheets in amyloid beta aggregates with an absorption band at ∼1614cm^-1^; so the possibility of certain beta sheet conformations exhibiting absorptions at lower than expected wavenumbers cannot be ruled out. Taken together, the deconvolution results indicate that the structural heterogeneity observed in the fibrillar and prefibrillar aggregates is reflective of an interplay between parallel and antiparallel beta structures and beta turns/random coils. Spectra with a prominent peak at between 1630-1635cm^-1^ represent a structural distribution that contains more parallel beta sheet, while those containing a dominant peak between 1656-1660cm^-1^ represent conformations that lack significant beta structure. And lastly, spectra with a peak at 1668cm^-1^ or higher represent conformations with distinct antiparallel structure in addition to turns. In the conventional view of amyloid aggregation, oligomers are believed to contain antiparallel beta sheets, which evolve into fibrils with parallel beta sheet structure. Our results show that oligomers generated in early stages of aggregation have significant structural variation, and the antiparallel beta sheet structure in oligomers becomes dominant with time. Given that protofibrils and fibrils exhibit the same structural heterogeneity as the early oligomers, and the abundance of ‘homogeneous’ oligomers alongside fibrils when they are expected to be incorporated into fibrils, these oligomeric species likely represent aggregation states that are less likely to be incorporated into fibrils. In that event, the heterogeneity can be argued to be favorable for formation of mature, ordered fibrils. Ahmad et. al. have shown that the structural transition to fibrils proceeds through oligomeric species that lack distinct beta structure, which is consistent with our observations, and support the hypothesis that the mature oligomers represent a conformation hindered in their ability to participate in fibril formation^15^. The observation of antiparallel beta sheet character in fibrils is somewhat unexpected and deviates from the commonly expected parallel beta sheet structure of mature fibrils. However, antiparallel beta sheet structure in fibrils has been reported for certain mutants of beta amyloid^45^ and also for ganglioside induced aggregation^46^. Our results show that early-stage fibrils of wild-type Aβ formed without the interference of any external agents can have some antiparallel character which possibly evolves into parallel structures upon maturation. It should be noted that a comprehensive assessment of the structural distributions in different aggregation states and their evolution into mature fibrils with parallel cross beta structure requires structural analysis of aggregates at many more time points, which we aim to address in future work.

**Figure 4.**
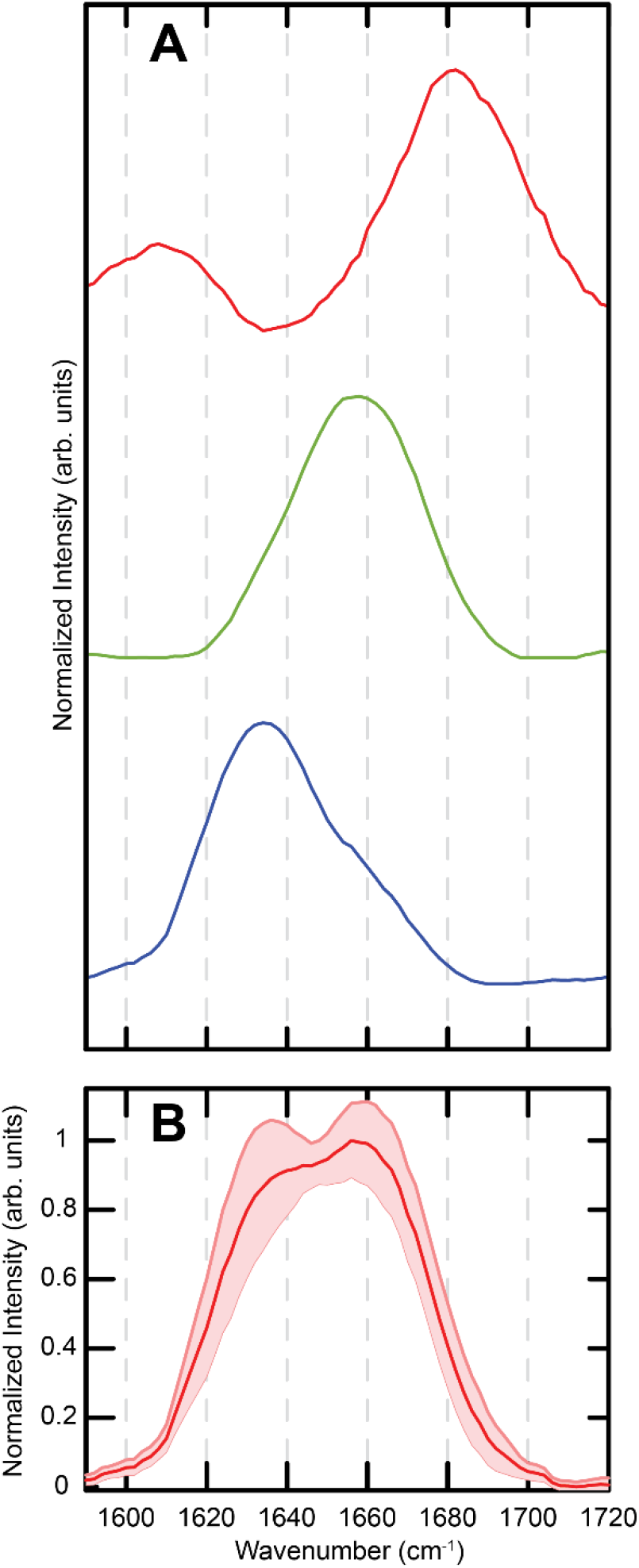
NMF deconvolution of IR spectra of aggregates. (A) Spectral components derived from NMF factorization of spectra. (B) Average spectrum of fibril showed in Figure 2A, determined by taking of spectra acquired along the fibril axis.

### Structural heterogeneity of fibrils is manifested in tissues

Observation of the aforementioned spectral heterogeneities leads rise to an important question, namely: do these persist in AD plaques? The spectral variations within a single fibril indicate that the average spectrum of individual fibrillar aggregates, shown in Fig. 4B, can be very different from the conventional infrared absorptions associated with parallel beta sheets owing to the inherent structural variations described earlier. AFM is not a technique well-suited for tissue imaging, and hence extension of AFM-IR towards characterizing plaques is experimentally challenging. Infrared confocal imaging, which integrates infrared spectroscopy with optical microscopy, is an approach ideal for tissue applications and allows for mapping the chemistry of amyloid aggregates in AD tissues while providing the same molecular contrast^47-49^. We have recently demonstrated the capabilities of infrared imaging towards identifying the chemical heterogeneities of plaques^47^, wherein we demonstrated that amyloid plaques of the same morphology can vary significantly in terms of their beta sheet content. The heterogeneity in the amyloid aggregates observed in this study are in excellent agreement with our prior work and provide a molecular-level explanation of the plaque chemical variations. To further validate and test the generality of our prior results, here we expand on this work and apply infrared imaging to map plaques in AD temporal lobe tissue. Fig. 5A-B show optical images of two immunohistochemically (IHC) stained cored plaques which are expected to contain fibrillar amyloid aggregates at their core. For visualization of the beta sheet content of the plaques, we plot the spatial map of the ratio of the spectral intensity at 1628cm^-1^ normalized to the intensity at 1660cm^-1^ in Fig. 5C-D, which has been demonstrated to be reflective of the beta sheet content in amide-I spectra^47, 49^. It can be clearly seen that the plaques, while expected to contain fibrils with parallel beta sheet structure, exhibit very little beta sheet signature as evidenced by lack of bright pixels within the plaque boundary. This is not representative of all amyloid plaques, and we also observe cored plaques with the expected beta sheet signatures (Suppl Fig. 6). A total of 19 cored plaques were identified through immunostaining in these tissues, and 8 contained beta sheets while 11 did not exhibit significant beta sheet signatures. The full tissue IHC and infrared ratio images are shown in Suppl Fig. 7 and 8 respectively. These observations are consistent with our prior reports and provide unequivocal evidence that the heterogeneities observed in individual amyloid aggregates can also manifest themselves in AD plaques and can thus also have potential clinical significance. The implications of these heterogeneities in AD progression and their prevalence in different stages of AD will be addressed in future work.

**Figure 5.**
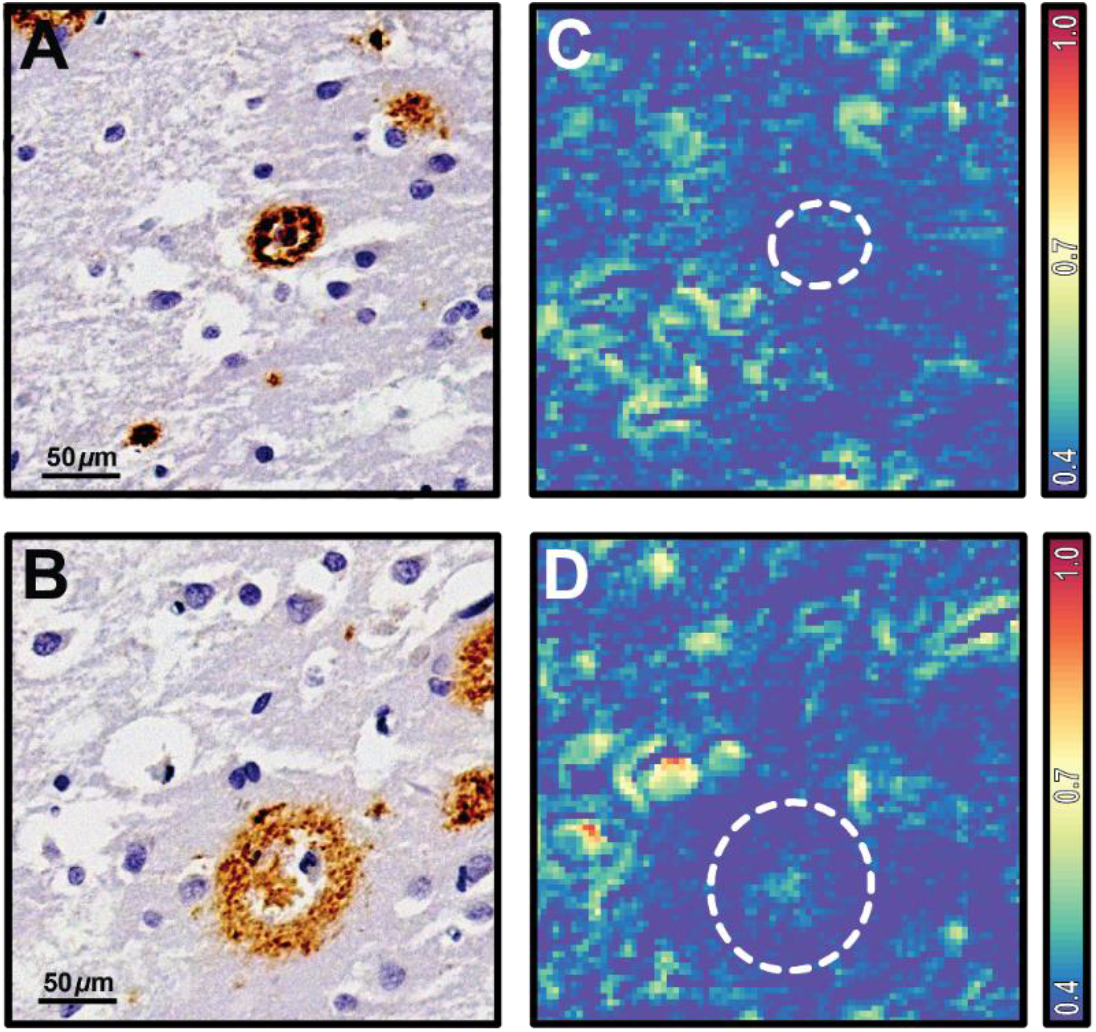
Infrared microscopy of amyloid plaques. (A-B) Optical images of immunohistochemically stained cored amyloid plaques. (B) Corresponding IR ratio images (1628cm^-1^:1660cm^-1^) depicting the abundance of beta sheets. The data were acquired from a diseased AD temporal lobe tissue section.

## Discussion

In this study, we have demonstrated for the very first time that early-stage amyloid aggregates are structurally heterogeneous, and these heterogeneities are transmitted from prefibrillar aggregates to fibrils. The heterogeneity is reflected in the infrared spectra and variations therein of individual aggregates. The structural variation along a single fibril species indicate that amyloid fibrils may not necessarily be perfectly ordered which is not apparent from spatially averaged measurements. Deconvolution of the spectra show that the distributions are rooted in the relative distribution of beta sheets and beta turns. These results point to a scenario where the average spectrum of fibrils can lack the typical beta sheet signatures, indicating that amyloid aggregates and beta sheets are not always synonymous as they are often interpreted to be. We have validated this observation through infrared microscopy of amyloid plaques in tissues and show that plaques that are expected to contain fibrils can often lack beta sheet structure.

In summary, the results reported herein underscore the need for spatially resolved spectroscopy of amyloid aggregates and strongly establish the capabilities of AFM-IR as a method of choice for identifying the structural variations within different subgroups of aggregates and their evolution. These results can pave the way towards subtyping of plaques using amyloid secondary structure and can potentially unlock novel therapeutic strategies for Alzheimer’s disease.

## Methods

### Preparation of Aβ42 aggregates

Aβ42 (rPeptide, USA) was treated with 1,1,1,3,3,3-hexafluoroisopropanol (HFIP) to remove any preformed aggregates. HFIP was evaporated under vacuum and the stock solution was prepared in 1X PBS buffer at pH 7.4. The aggregation was carried out at 37ºC without any agitation.

### Preparation of samples for AFM-IR experiment

Samples for AFM-IR experiments were prepared on ultraflat gold surface (Platypus Technologies, USA). 5 µl of the aggregation mixture was deposited at the center of the gold surface and incubated in a covered petri dish for 5 min. The substrate was subsequently gently rinsed with 500 µl milli-q water and dried under a stream of nitrogen gas.

### AFM-IR experiment

AFM-IR experiments were performed with Bruker NanoIR3 equipment having mid-IR Quantum Cascade Laser (MIRcat, Daylight solutions). All AFM experiments were carried out in room temperature in tapping mode. The instrument was continuously purged with dry air to keep the relative humidity of less than 5% during the data collection. The cantilever resonance frequency has been 75 ±15 kHz and the spring constant is 1-7 N/m. Scan speeds were varied from 0.5-1.0 Hz. First a high-resolution AFM image of the sample were captured and then IR spectra has been recorded at different regions of interest. Spectra were recorded with 2 cm^-1^ resolution. 128 co-additions for each spectral point and 16 co-averages were applied for each spectrum.

### Tissue imaging with infrared microscopy

Formalin fixed paraffin embedded (FFPE) human temporal lobe diseased tissue was purchased from Biochain Institute. The postmortem tissue specimens studied in this work are deidentified and were determined to be not human subjects research by the Office of Research Compliance at the University of Alabama. These tissues were deparaffinized in hexane for 24 prior to imaging. The microscope design is based upon work published by Bhargava and coworkers^50^, and has been described in detail elsewhere^47, 50^. The microscope uses a tunable Quantum Cascade Laser (Block Engineering) as the illumination source and a TE cooled mercury cadmium telluride (MCT) detector (Boston Electronics). All images were acquired using a 0.74NA objective. Infrared absorption images at discrete wavenumbers of interest were acquired with a pixel size of 2 microns. All measurements were made in transflection mode on infrared reflective low-emissivity slides (Kevley Technologies). The discrete frequency images were aligned using the image processing toolbox in MATLAB prior to ratioing to minimize any scanning artifacts. Adjacent tissue sections were stained with the anti-amyloid MOAB-2 antibody (Sigma Aldrich) at the University of Alabama Birmingham pathology core research lab. Brightfield optical images of stained tissues were acquired at 40x magnification using an Olympus BX43 microscope.

### Spectral analysis

Spectra were smoothed using (3, 7) Savitzky-Golay filter and a linear baseline correction was performed. All spectral analysis was carried using the MATLAB software package.

### Spectral deconvolution using Non-Negative Matrix Factorization

NMF is a blind spectral deconvolution approach which factors the spectra into a weighted sum of components or sub-bands similar to singular value decomposition (SVD), which has been used successfully for deconvolution of infrared spectra^40-42, 51^. The key difference between NMF and SVD is that the latter allows for spectral components to have negative values, whereas NMF does not. Since infrared absorbances cannot be negative, NMF is a more suitable choice for infrared spectral deconvolution that produces physical intuitive results. In NMF, the non-negative spectra matrix S into decomposed into two lower -ranked non-negative matrices as: S = M·C. Here M is a matrix that contains the component spectra, while C contains the weights or populations of each component. The optimal matrices M and C are iteratively determined to minimize the residual between the original spectral data the product of the lower rank approximations. The NMF analysis was performed on the entire spectral set, which makes this approach similar to global fitting: essentially every spectrum is decomposed as a weighted sum of the same components. The number of components was determined to be 3 using the L-curve approach^52-53^.

## Acknowledgment

This work was supported by the National Institutes of Health (Award 1 R35 GM138162 to A.G.). The authors thank Dr. Dezhi Wang at UAB Pathology Core Facility for help with IHC staining.

## Materials and Correspondence

Correspondence should be addressed to Ayanjeet Ghosh

## Author Contributions

S.B. prepared the amyloid aggregate samples and performed the AFM-IR experiments and data analysis. B.H. performed the infrared imaging experiments and data analysis. S.R. performed spectral deconvolution. A.F. acquired optical images of stained tissue and spatially located plaques. S.B. and A.G. conceived and designed the study and wrote the manuscript.

## Supplementary Information

AFM image of oligomers and their height distribution, additional IR spectra for oligomers, protofibrils and fibrillar aggregates; AFM image and height distribution of protofibrils and fibrils; AFM image of oligomers present in the fibril sample; infrared images of amyloid plaques exhibiting beta sheet signatures.

## Data Availability

All data needed to evaluate the conclusions drawn in this communication are included in the manuscript and the Supplementary Information file. The datasets and any other additional data are available from the corresponding author upon reasonable request.

