## Supplementary Information for "Structural heterogeneity of amyloid aggregates identified by spatially resolved nanoscale infrared spectroscopy"

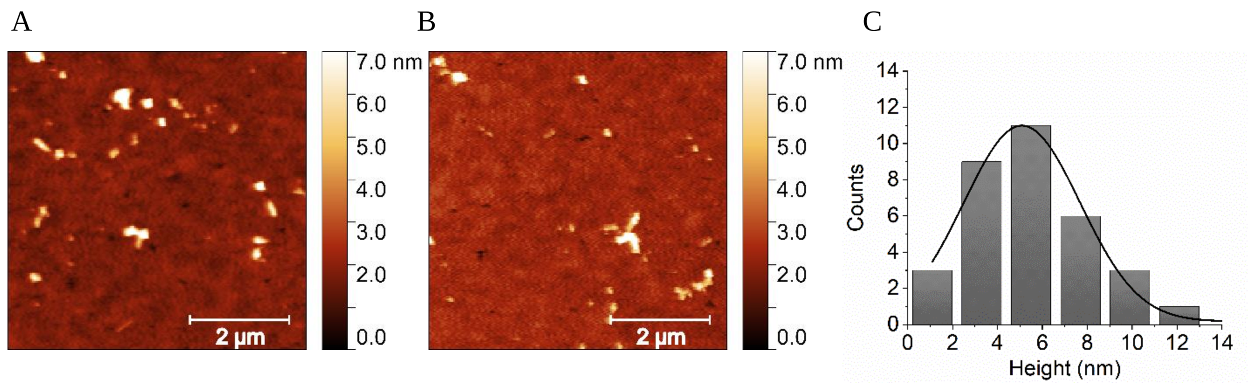


**Supplementary Figure 1.** (A-B) AFM height image of Aβ42 oligomers observed at two different areas on gold substrate. Primarily oligomers were distributed as isolated species allowing spectroscopic measurements from individual oligomers. (C) The histogram shows the height distribution of the oligomers.

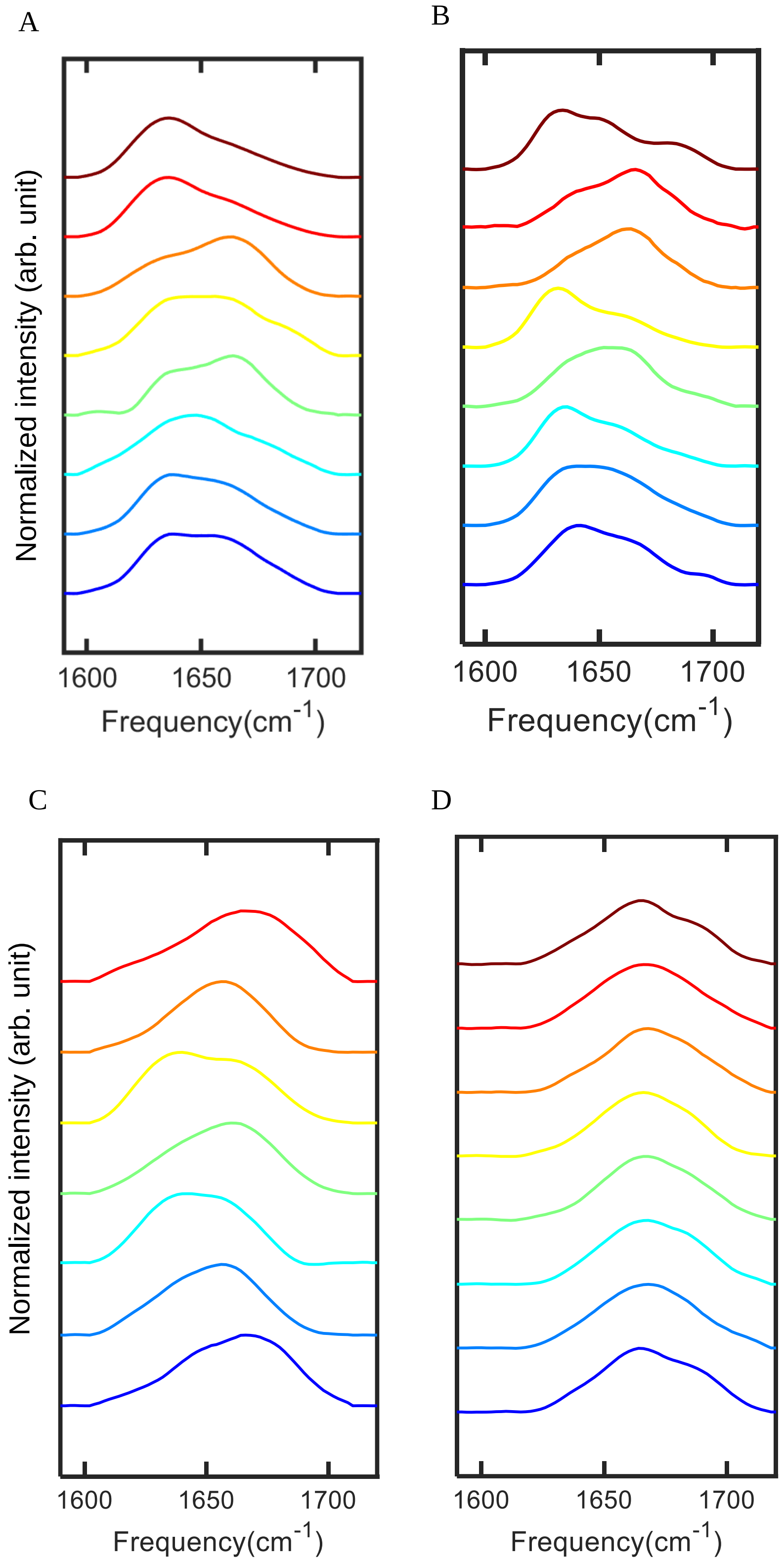


**Supplementary Figure 2.** Spectral heterogeneity of amide I band region of the IR spectra recorded on (A) Aβ42 oligomers, (B) protofibrillar aggregates and (C) fibrils. (D) Spectral heterogeneity is not evident in the oligomers which are present with fibrils.

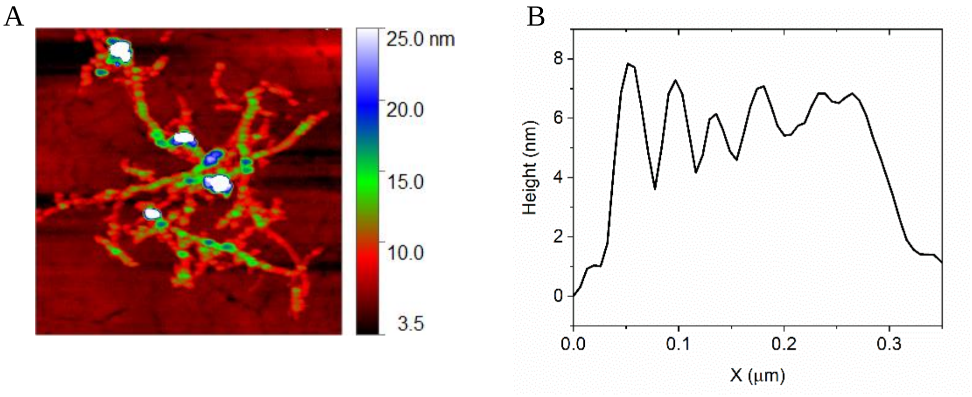


**Supplementary Figure 3.** (A) AFM topographic image of Aβ42 protofibrils shown in rainbow height scale. The presence of different colors on the protofibrils depicts the periodic changes in the height value, indicating a non-smooth, beads like conformations composing the protofibrillar aggregates. (B) A line-profile obtained from the image shown in A, along the long axis of protofibril. The corrugated profile also indicates the presence of globular features as an essential constituent of protofibrils.

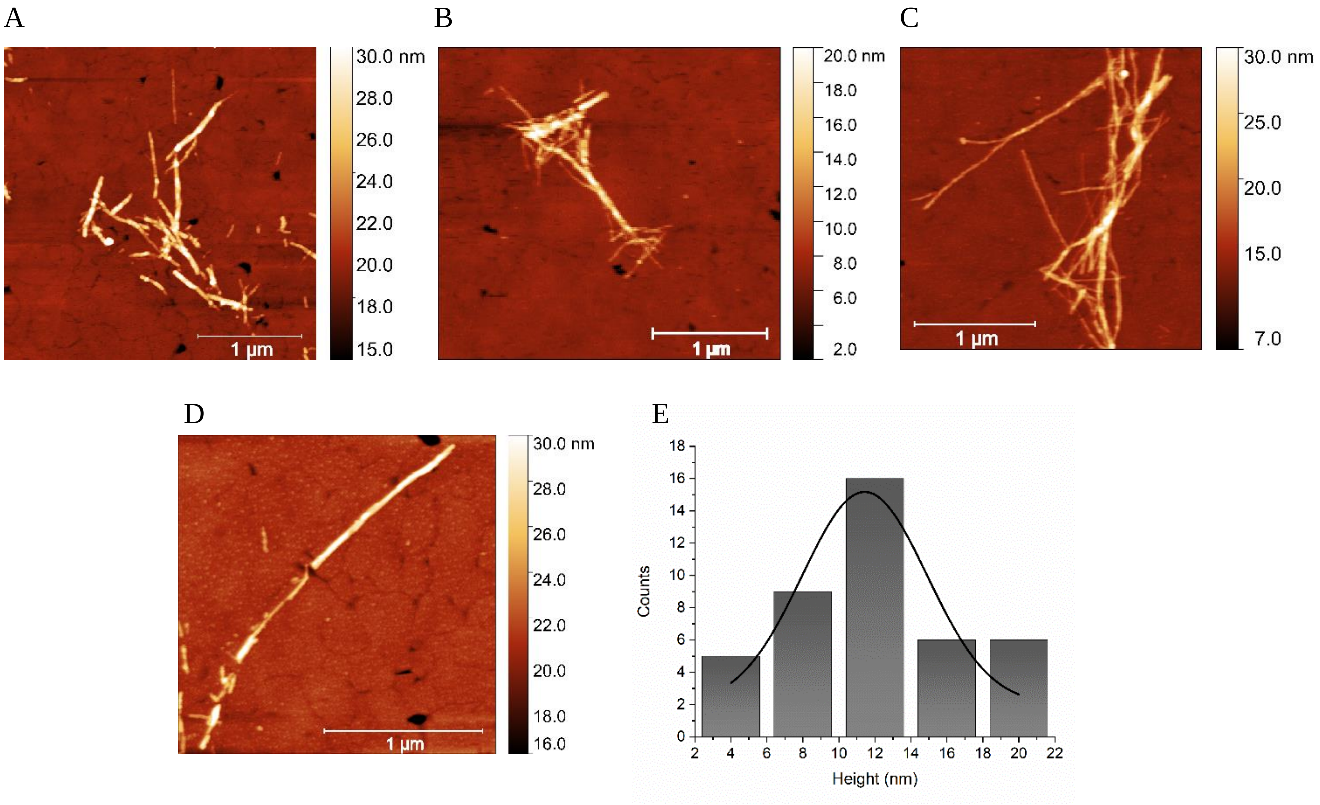


**Supplementary Figure 4.** (A-B) AFM topographic image of Aβ42 fibrils at two different areas of the surface. Fibrils are relatively smooth, where the beads like structure, observed in protofibrils, is not clearly evident. (C) The histogram shows the height distribution of fibrils.

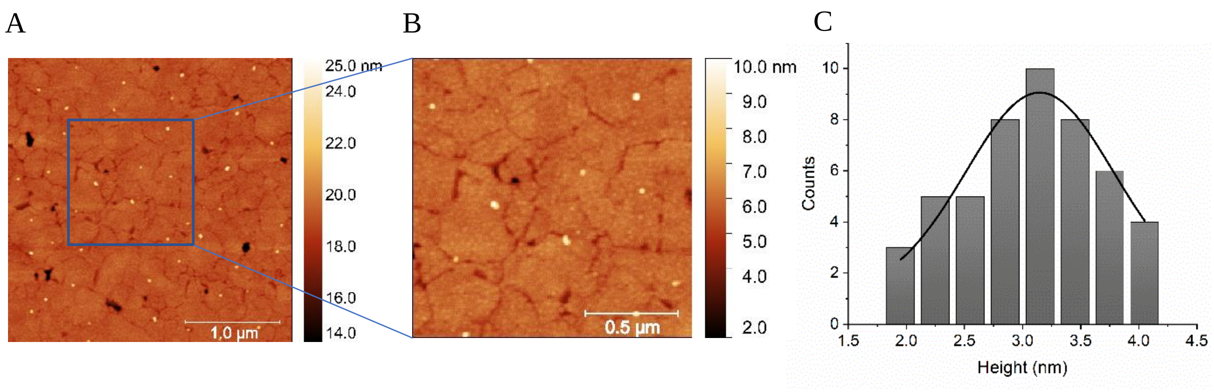


**Supplementary Figure 5**. (A) AFM topographic image of Aβ42 oligomers which are present in the same sample with fibrils. (B) Zoomed image of the area shown in A. Oligomers are primarily globular in shape. (C) Height distribution of the oligomers.

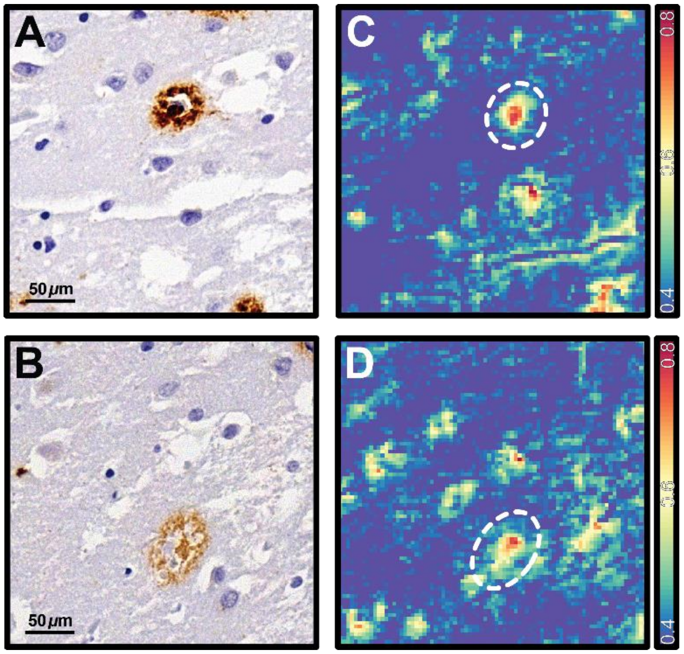


**Supplementary Figure 6**. (A-B) Immunohistochemically stained amyloid plaques from AD temporal lobe tissue. (C-D) Corresponding infrared ratio images (1628cm$^{-1}$:1660cm$^{-1}$).

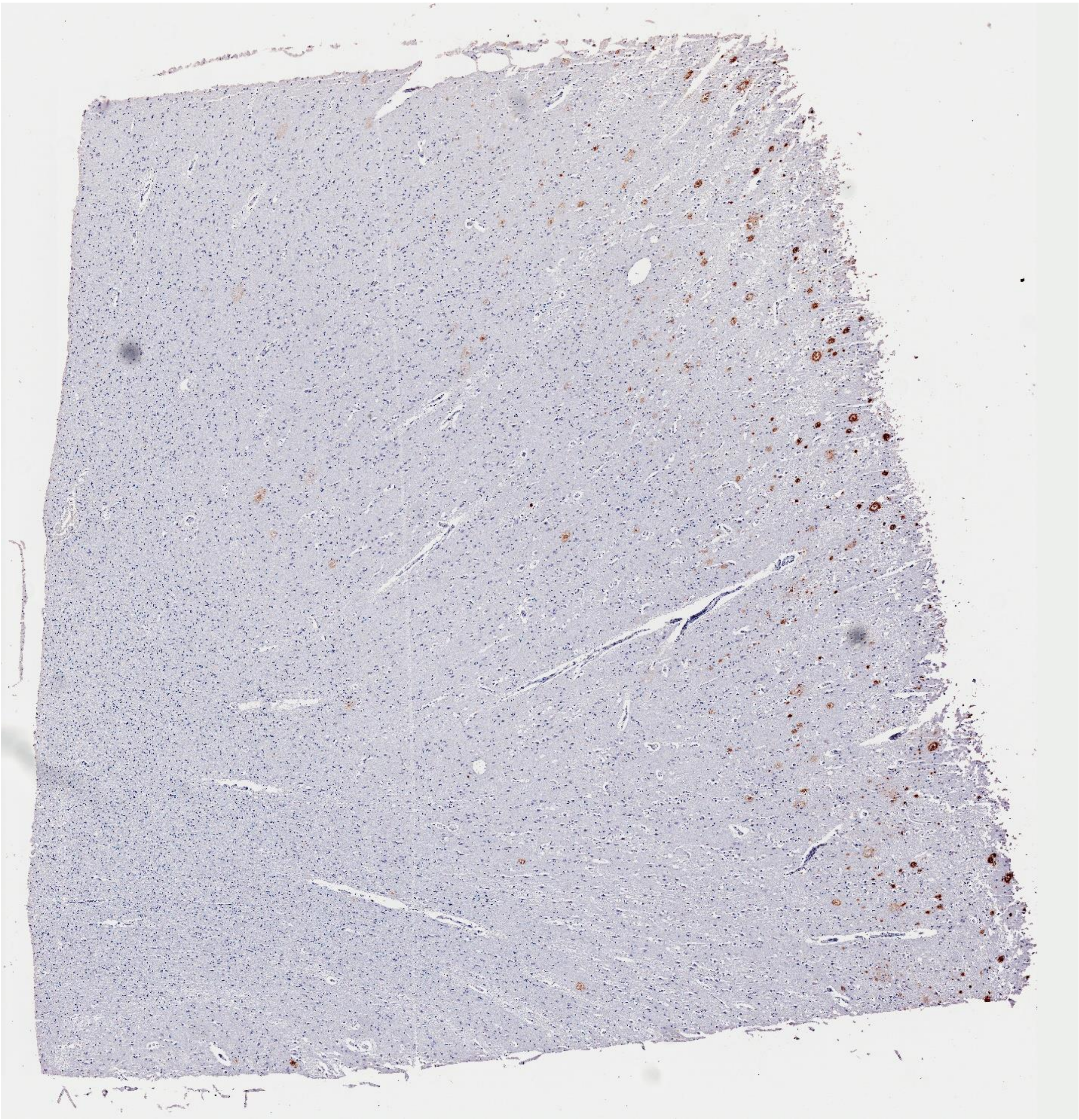

**Supplementary Figure 7**. Optical image of immunohistochemically stained entire AD temporal lobe tissue.

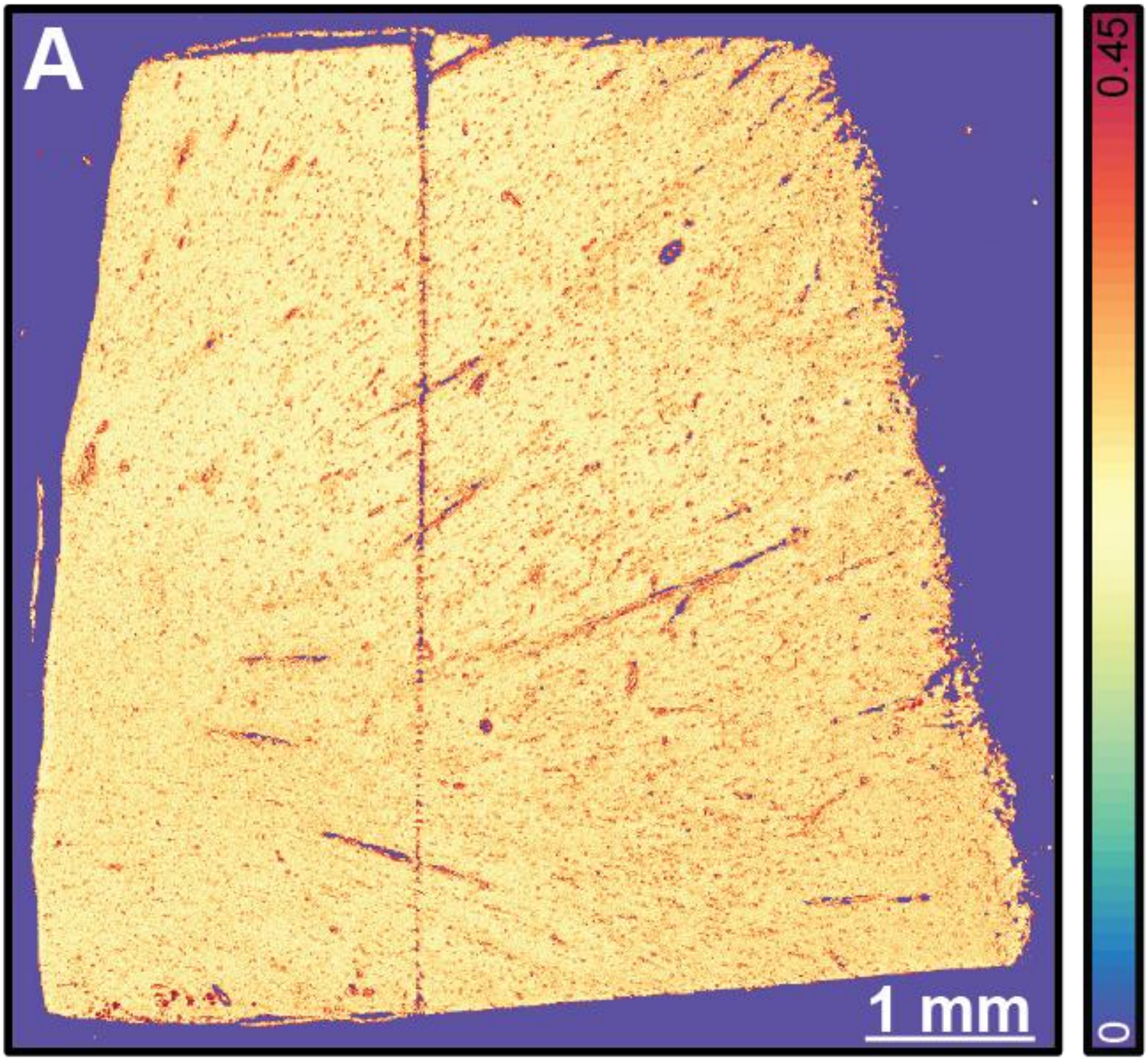


**Supplementary Figure 8**. Infrared ratio image (1628cm$^{-1}$:1660cm$^{-1}$) of entire AD temporal lobe tissue.
